# Light pollution affects West Nile virus exposure risk across Florida

**DOI:** 10.1101/2020.05.08.082974

**Authors:** Meredith E. Kernbach, Lynn B. Martin, Thomas R. Unnasch, Richard J. Hall, Rays H.Y. Jiang, Clinton D. Francis

## Abstract

Emerging infectious diseases (EIDs), including zoonotic arboviruses, present a global health threat. Multiple components of human land use change have been linked to arboviral emergence, but one pervasive factor that has received comparatively little attention is light pollution. Although often considered a component of built environments, artificial light at night (ALAN) outpaces the growth and spatial extent of urbanization, and thus affects areas where human population density and anthropogenic land changes are modest. West Nile virus (WNV) emergence has been described as peri-urban, but recent research suggests that its relative ubiquity in human-altered environments might actually be due to ALAN. Indeed, we found previously that experimental ALAN exposure enhanced avian competence to transmit WNV to mosquitoes. In the present study, we asked whether such organismal effects manifest ecologically by determining whether WNV exposure among sentinel chickens in Florida is related to local ALAN conditions. We found strong support for a nonlinear relationship between ALAN and WNV exposure in chickens with peak WNV risk occurring at low ALAN levels. Importantly, effects of ALAN on WNV exposure were stronger than other aspects of urbanization; only ambient temperature in the month prior to sampling had a comparable effect to ALAN. These results represent the first field evidence that ALAN might affect infectious disease exposure risk. We advocate for further research on how ALAN influences zoonotic risk, as well as efforts to study alternative nighttime lighting methods to reduce such risk.

**Significance Statement:** Light pollution associated with human development is a globally pervasive and rapidly expanding anthropogenic stressor; but despite documented effects on host immune functions and vector behaviors, how it affects infectious disease risk is unknown. Using data from the Florida Department of Health arbovirus surveillance program, we show that light pollution is a stronger predictor of variation in West Nile virus (WNV) exposure risk than many other previously implicated anthropogenic and natural environmental variables. Light pollution effects are nonlinear, so risk is highest in areas with dim light pollution. Our results highlight a new way that light pollution might affect human and wildlife health.

## Introduction

Emerging infectious diseases (EIDs) are among the greatest threats to public health today (1, 2). Most EIDs are zoonotic in origin (70%), in that causative agents spill over to human populations from other species (3, 4). Anthropogenic effects on wildlife can thus become detrimental to humans in places where humans and wildlife come into contact. The recent surge in many EIDs can be attributed to various forms of global change including alterations of the climate and the structure and biological composition of landscapes (3, 5, 6). One of the more recent examples of an anthropogenically-driven zoonotic EID is West Nile virus (WNV), which was introduced to the United States in 1999 (7). WNV decimated songbird populations within the first several years of its arrival, and its propensity to be transmitted by so many vector species also made it a source of substantial human disease. Now, 20 years since its introduction, WNV continues to cause harm to human and avian populations, particularly in highly modified areas. Some factors that influence WNV transmission (e.g., weather-driven mosquito population growth) have been identified; however, we still lack a thorough understanding of other potential drivers (8).

WNV is recognized as a peri-urban arbovirus, as human incidence and songbird seroprevalence is much higher in or near urban habitats (9, 10). Historically, a higher incidence of WNV in or near cities was linked to aspects of environments that influence mosquito success such as local climate and availability of breeding sites (10, 11). The Florida Department of Health surveillance system has closely monitored arbovirus transmission in mosquito breeding hotspots, such as peri-urban drainage systems, since the late 20^th^ century (8, 12). WNV dynamics resemble other zoonoses in that urban and agricultural development affect emergence and transmission (13). For instance, Lyme disease (caused by *Borrelia burgdorferi*) and flavivirus infections including yellow fever, dengue, and chikungunya, all emerge in developed areas where biodiversity is often low, host density is high, and vectors thrive in close proximity to humans (14–16). Some anthropogenic stressors have been found to affect risk, but many conspicuous and common ones have never been considered, including light pollution. One form of light pollution, artificial light at night (ALAN), covers 18.7% of the continental U.S. and affects 99% of the human population with increases anticipated in the future [e.g., 2.2% increase per year from 2012 to 2016, (17)]. Further, small areas of urban development such as city centers can emit light into distant suburban and rural landscapes, suggesting that light pollution effects may be widespread (18, 19).

Light pollution also affects multiple host and vector traits with consequences for disease transmission. For instance, in night shift workers, ALAN can affect non-communicable disease risk (e.g., cancer, diabetes, etc.) (20). ALAN also induces dysregulation of immune responses in lab rodents, likely because vertebrate immune systems are ‘fundamentally circadian in nature’ (20, 21). House sparrows, an urban-residing passerine reservoir of WNV, experimentally infected with WNV and exposed to modest ALAN (i.e., 5 lux; a full moon on a clear night is 0.3 lux) maintained transmissible WNV titers for 2 days longer than controls but did not incur higher mortality (22, 23). Epidemiologically, these effects were estimated to increase outbreak potential by 41% (22). Furthermore, arthropod vectors of WNV are renowned for their flight-to-light behaviors; for those that survive desiccation or depredation, flight-to-light behavior might concentrate infection risk where light pollution is common (24, 25). WNV can be transmitted by as many as 45 vector species, many of which bite wildlife (preferentially songbirds) and humans (26, 27). Given the current pervasiveness of ALAN, we asked whether light pollution can affect infectious disease dynamics in a part of the US where arboviruses and common and influential, both economically and socially (28).

Specifically, we investigated whether ALAN affected risk of WNV exposure across several counties of Florida where emergence and spillover has occurred in the recent past (29). Although several forms of light pollution could impact vector borne infections, we focused on WNV because it is the most broadly distributed arbovirus and most important causative agent of viral encephalitis worldwide (30, 31). Using data from the Florida Department of Health (FDOH) sentinel chicken WNV surveillance program (12), we tested the hypothesis that WNV exposure, measured as the number of sentinel chickens undergoing antibody seroconversion, would be highest at low to mid ALAN levels through the potential combined effects of increased vector density (via ALAN attraction), vector-host interactions and subsequent biting rates, and/or extended host infectious periods (20, 24). We also hypothesized that WNV exposure might be lowest in extreme, or intense, ALAN conditions where host and vector predation and pathogen-induced mortality risk might be high (32, 33). We used mixed-effect models with and without spatial correlation structure to assess the effects of ALAN on WNV exposure for four recent years across 5 counties totaling over 1000 events from 100 unique geographical coordinates, including many peri-urban regions (Figure 1). These models also accounted for previously documented and other hypothesized predictors of WNV risk (i.e. weather, soil moisture, and several aspects of urbanization).

**Figure 1.**
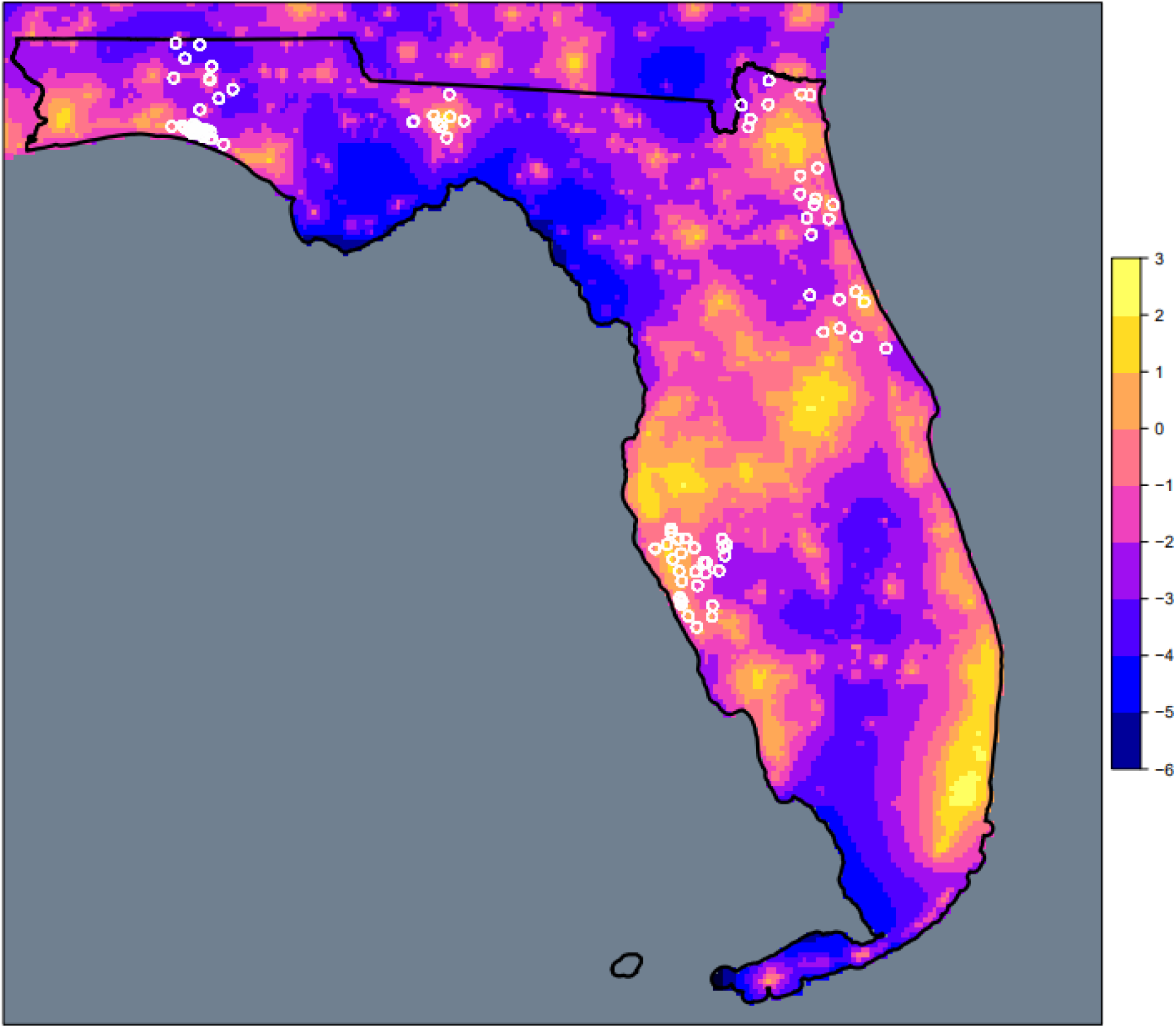
Sentinel chicken sampling locations (white circles) throughout Florida overlaid on radiance [mcd/m^2 (millicandela/m^2)] estimates from Falchi et al. (56). Color ramp depicts light pollution intensity in units of log radiance.

## Results and Discussion

We found that ALAN was a strong but nonlinear predictor of WNV exposure risk in Florida (Figure 2A). Models with and without spatial correlation structure (or an offset to account for variation in the number of chicken samples per site; see supplementary text) were equally competitive (Table 1) and included qualitatively similar parameter estimates for ALAN. Specifically, we found that at low ALAN levels (i.e., at radiance intensities less than 1), WNV risk was greatest, but risk declined towards intermediately polluted sites (Figure 3A), which was consistent with our hypothesis.

**Table 1.** WNV exposure model selection including spatial, non-spatial, and offset options. Rankings among models with and without spatial correlation structure and offsets to account for variation in number of sentinel chickens surveyed.

| Model | logLik | AIC | ΔAIC |
| --- | --- | --- | --- |
| Spatial | -727.492 | 1492.984 | 0.000 |
| Non spatial with offset | -729.939 | 1493.878 | 0.895 |
| Non spatial | -729.800 | 1493.601 | 0.617 |
| Spatial with offset | -729.272 | 1496.543 | 3.560 |

**Figure 2.**
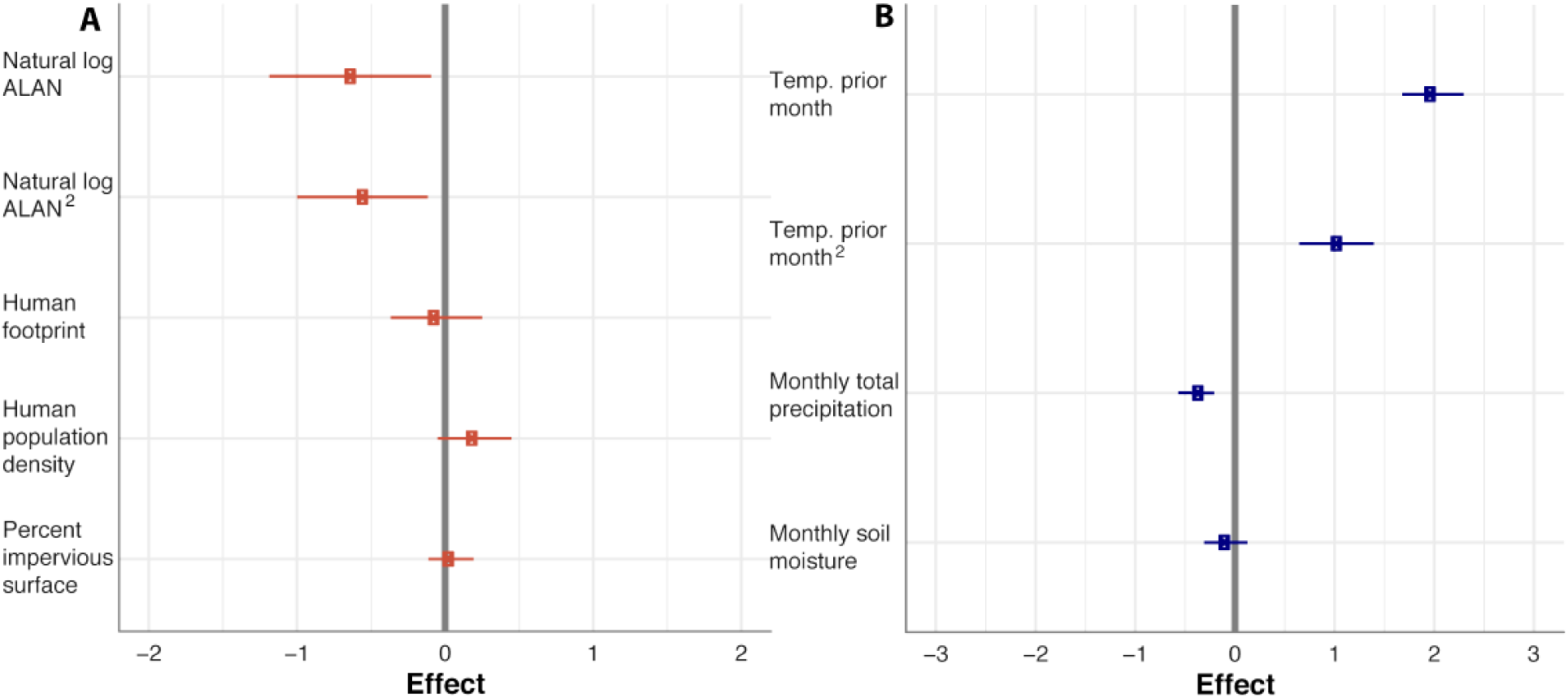
Effect sizes of (A) ALAN and urbanization, and (B) temperature and weather variables on West Nile virus risk. Standardized effect sizes (square points) and 95% CI (lines) for (A) anthropogenic and (B) weather-related variables from top ranked model in Table 1. (A) Natural log of ALAN and ALAN^2^ in radiance [mcd/m^2 (millicandela/m^2)] had the largest and only strong effect sizes compared to urbanization parameters, whereas (B) mean temperature and mean temperature^2^ of the prior month had the largest effect sizes compared to precipitation and soil moisture variables. Variables are natural log transformed to account for the extreme values in the distribution of data.

**Figure 3.**
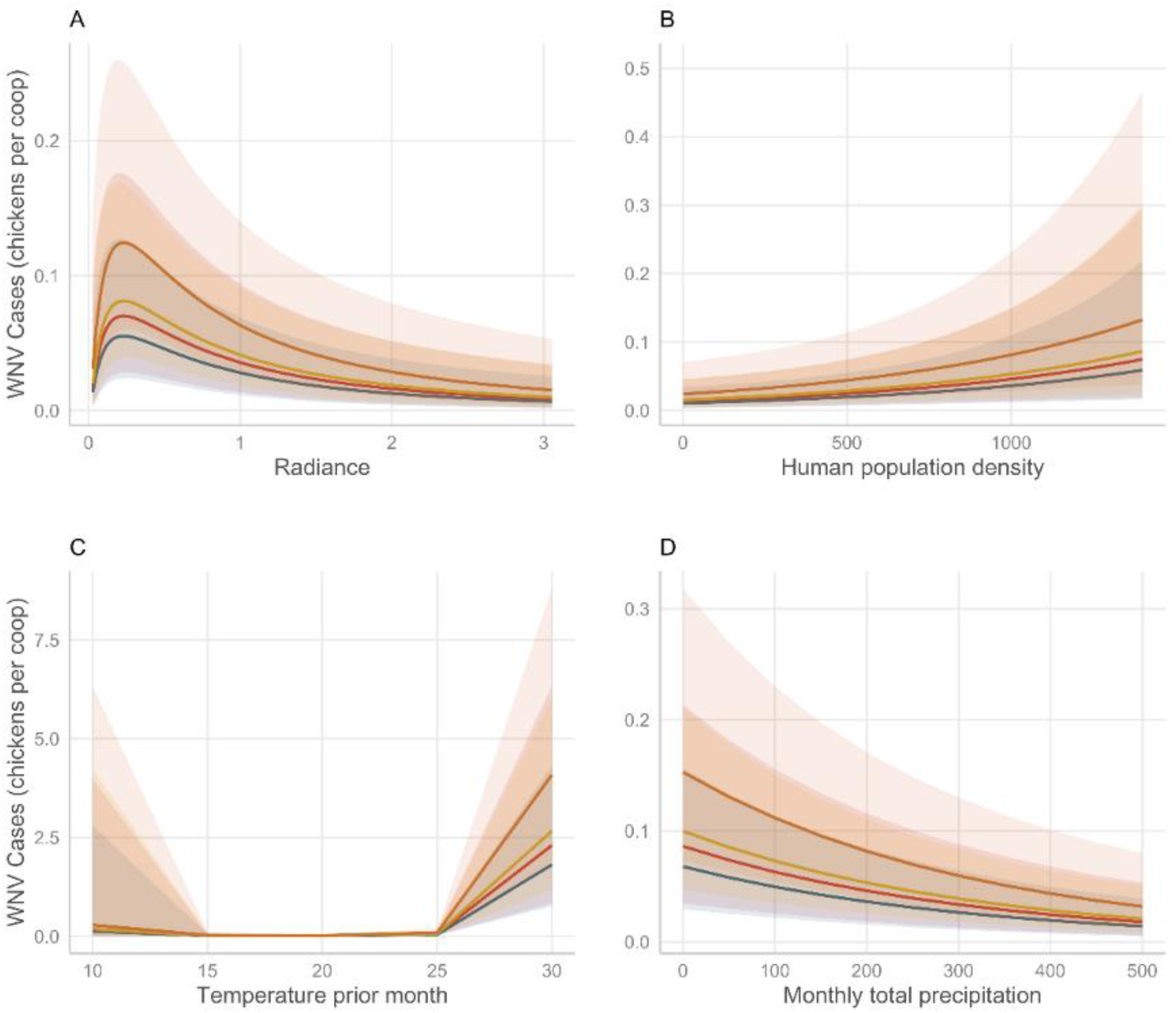
Interannual effects of ALAN (A), human population density per km^2^ (B), mean temperature of the previous month in degrees Celsius (C) and monthly total precipitation (mm) on incidence of WNV seroprevalence (D). Effects of each plotted by year with 95% CI band: red = 2015, blue = 2016, yellow =2017, orange = 2018. Effects of ALAN, human population density, temperature, and precipitation on WNV exposure were consistent across 5 years.

Besides the consistent influence of ALAN, all models provided strong evidence for a lag effect whereby high temperature (>25 °C) during the previous month strongly and positively predicted WNV seroprevalence (Figure 2B, Figure 3C). In contrast, increases in monthly cumulative precipitation was negatively related to WNV seroprevalence (Figure 3D), which is also consistent with prior work (34). In contrast with previous reports (9, 10), we found no influence of two prominent metrics of urbanization, human footprint index or percent anthropogenic impervious surface, on WNV seroprevalence. We found mixed support for an influence of human population density on WNV seroprevalence where two of the four models suggested there was a small positive relationship (Figure 3B; supplementary text). Additionally, there was also some evidence for interannual variation in WNV case incidence; nonetheless, we found consistent ALAN effects on WNV risk across several years (Figure 3A). Finally, to ensure that the relationship between WNV seroprevalence and ALAN did not reflect nonlinear effects of urbanization in general, we also tested for polynomial effects of human population density, human footprint index, and percent anthropogenic impervious surface. No polynomial terms for any of these predictors were supported (Table 2). To our knowledge, ours is the first study to indicate that light pollution influences arbovirus infection risk under natural conditions. Our results also hint that some previous observations for elevated arboviral risk in peri-urban areas might have partly been indications of light pollution effects. Below, we discuss how light pollution might exacerbate risk of WNV and other EIDs, and why ALAN had a nonlinear effect on WNV risk.

**Table 2.** Sensitivity analysis of polynomial effects on WNV exposure. Rankings among models where anthropogenic predictors and mean temperature of the previous month were modeled with linear or polynomial effects. All predictors remained in each iteration of the model and tested polynomial effects (i.e., “poly”) are listed prior to the parameter under consideration. * reflects top ranked model from Table 1. ALAN = artificial light at night, temp = mean temperature of the previous month, imp sur = percent anthropogenic impervious surface, pop den = human population density and hum foot = human footprint index.

| Model | logLik | AIC | ΔAIC |
| --- | --- | --- | --- |
| Spatial: poly ALAN, poly temp | -727.492 | 1492.984 | 0.000 |
| Spatial: poly temp | -730.790 | 1497.580 | 4.596 |
| Spatial: poly pop den; poly temp | -730.753 | 1499.506 | 6.522 |
| Spatial: poly imp sur; poly temp | -730.754 | 1499.508 | 6.524 |
| Spatial: poly hum foot; poly temp | -730.781 | 1499.561 | 6.577 |
| Spatial: poly ALAN | -736.806 | 1509.611 | 16.627 |
| Spatial: all linear | -740.050 | 1514.100 | 21.116 |

### Why does ALAN affect WNV exposure?

Although not previously demonstrated in natural conditions, the relationship between light pollution and WNV exposure was not altogether surprising. Recent experimental work demonstrated that ALAN exposure can extend the infectious period of one host, which, if such effects persist in nature, could increase local WNV risk (as quantified by the basic reproductive number, R_0_) by 41% (28). Other literature strongly suggests that ALAN could also increase the local density and feeding period of crepuscular vectors, which might significantly alter opportunities for transmission, as bite rate (compared to other terms) disproportionately affects WNV outbreak potential (35). Such effects are particularly likely because many vectors exhibit flight-to-light behavior, which could concentrate risk spatiotemporally (25). Transmission risk may be further exacerbated if short-wavelength light is present. This light form is used to census *Culex* mosquitoes, one of the many competent vectors of WNV. In the context of ALAN, risk might be elevated when light pollution is intense enough, or if the appropriate color temperature, to elevate vector densities (36).

### Weather, precipitation, and WNV

In addition to ALAN, average temperature of the prior month predicted WNV exposure in parts of Florida. It is unsurprising that this temperature measurement was a vital component in our models, as many aspects of WNV transmission are temperature-dependent (e.g., vector development, survival, and competence) (37). Indeed, most vectors of WNV thrive when temperatures increase over summer months, as high temperatures accelerate vector growth rates and extrinsic incubation periods (i.e., the time required to develop transmissible virus in salivary glands) (38). Periods of heavy rainfall and the availability of water sources for breeding can also sustain large populations of *Culex nigripalpus*, a moderately competent WNV vector in south Florida (39, 40). Conversely, severe drought spells are related to higher WNV incidence, which may reflect changes in host competence rather than vector success (34). Even after climate and weather variables were accounted for, substantial spatial heterogeneity in WNV incidence remained unexplained.

### Urbanization and zoonotic exposure risk

Urbanization has long been viewed as a driver of WNV emergence (10), but here, multiple metrics of urbanization had no to little predictive power for WNV risk. Other studies have concluded that urban land use and human population density were important predictors of inter-annual WNV prevalence over large geographic regions (13, 41, 42). These environmental features are hypothesized to drive WNV incidence due to decreased host diversity in urban areas, altered vector ecology (e.g., small water sources ideal for mosquito breeding), and/or increased host susceptibility to infection (43, 44). However, besides modest support for human population density, we found minimal effects of urbanization on risk. One potential reason we did not detect such effects might have been because of a lack of data from the most urbanized areas in Florida. Such data were unavailable from city centers simply because sentinel chickens are not housed there (supplementary text). However, our study included locations where anthropogenic impervious surface was 67% and the human footprint index was 46.28 on scale of 0-50, so we are confident that our locations were sufficiently urbanized for comparisons of urbanization and ALAN (45).

### Nonlinearity of ALAN effects on WNV exposure risk

Our results suggest that although WNV exposure risk initially increased with ALAN at low intensities, it subsequently declined towards intermediate-intensity ALAN sites. This phenomenon could have been driven by higher infection-induced mortality rates in maintenance hosts (resulting in decreased infectious period) or avoidance of light at night (46). Additionally, risk of host and vector predation could be higher in brightly lit areas, which could deplete host and vector densities (47). Many characteristics of highly light polluted areas (e.g., fragmented habitat supporting fewer hosts, fewer vector breeding sources, extensive vector control efforts) could reduce WNV incidence (48–50). To fully understand the mechanisms underlying the nonlinear relationship between WNV exposure and ALAN, we advocate for research that investigates the effects of ALAN on key drivers of transmission, vector survival, host susceptibility (35).

### Conclusions and Implications

Our study suggests that light pollution might play an important role in the emergence of arboviruses, including WNV, in Florida and elsewhere vectored pathogens are common. Investigating how host and vector responses to ALAN exposure vary with spectral composition and intensity level of light will be vital for a full understanding of the patterns in WNV exposure risk observed here. Many cities and neighborhoods are switching to cool white light-emitting diodes (LEDs), which could be especially harmful given the sensitivity of wildlife and humans to blue wavelengths (51). One economical, environmentally friendly solution might include a change to long-wavelength nighttime lighting, which is already widely used along shorelines to prevent drawing marine turtle hatchlings inland (52). Indeed, some studies on various wildlife species have also confirmed that exposure to red or amber-hued light at night prevents immunological and behavioral dysregulation (51, 53). As LED lighting has started to replace high pressure sodium and halogen bulbs, this change in lighting practices could provide a unique opportunity to continue to use LEDs that offer high energy efficiency without posing health threats to the wildlife and human community. Since little is currently know about differential sensitivities of vertebrate and arthropod vision to distinct wavelengths of light, we recommend that mitigation options involving alterations to the timing of intensity of illumination should also be explored.

## Methods

### Sentinel Data

Sentinel chicken data were shared by the Florida Department of Health from Leon, Manatee, Nassau, Sarasota, St. Johns, Volusia, and Walton counties for the years 2015-2018. The data were organized as number of sentinel chickens per site that tested positive for WNV antibodies out of 6 (89.5% of observations) or 4 (10.5% observations) total chickens tested each month. Our count data reflect typical numbers of WNV seroconversion in sentinel chickens in Florida (29). Once a chicken tested positive for WNV antibodies, it was removed from a site and replaced with a WNV-naïve chicken, ensuring that all positives are new exposures. Coops housing sentinel chickens were occasionally moved small distances resulting in small differences in coop locations across years (typically < 0.001 degrees or less than approximately 100m, but as much as 11km). As such, we used the unique coordinates from each sentinel chicken sampling location to determine radiance (ALAN), weather variables (soil moisture, temperature, precipitation), and urbanization variables (human population density, anthropogenic impervious surface, human footprint index, see below).

### Environmental Data

Because multiple dimensions of the environment, including ALAN, vary along an urban gradient, understanding which aspects are responsible for elevated WNV risk is paramount. As such, we assembled geospatial data reflective of several dimensions of urbanization: percent anthropogenic impervious surface from the 2011 National Landcover Database [30m resolution;(54)], human population density from the 2010 US Census [1km resolution; (55)], human footprint index data reflective of conditions in 2009 [1km resolution; (45)], plus ALAN estimates from the world atlas of artificial night sky brightness [1km resolution each; (56)], which was derived from NASA’s VIIRS instrument. We temporally-harmonized high-resolution monthly cumulative precipitation and monthly mean temperature data (0.8km resolution) from the PRISM database (57) and monthly mean soil moisture estimates (3km resolution) generated from NASA’s Sentinel 1/SMAP platform (58) to match the month of sentinel chicken sampling. Additionally, to account for potential lags in conditions that could favor vector abundance, we also collected data for these variables from the month prior to sampling. Because >94% of the positive detections of WNV occurred after May of each year, we restricted our analyses to June-December. Additionally, because mean soil moisture estimates for the month of surveillance and the prior month were not available for all records, we restricted analyses to those records with complete environmental data, resulting in a final dataset of 1126 surveillance events from 80 sites and 105 unique coordinates.

### Data analysis

We modeled incidence of WNV seroprevalence using mixed-effect models with and without a spatially-explicit exponential correlation structure with negative binomial error and implemented in the R packages *spaMM* 2.7.5 (59) and *lme4* 1.1-21 (60). Given the hierarchical and repeated sampling regime of sentinel chicken surveillance programs, we included nested random effects of sampling month within site within county. For fixed effects, we built a model that included ALAN, the three variables reflective of urbanization (i.e., impervious surface, population density, human footprint) and monthly mean soil moisture, precipitation total, and mean temperature, plus year to account for interannual variation. All continuous variables were centered and scaled to facilitate direct comparisons of effects and, based on our hypothesis that WNV seroprevalence would peak at intermediate ALAN exposure levels, we modeled the effect of ALAN as a second-order polynomial. Preliminary data exploration revealed a nonlinear relationship between WNV seroprevalence and monthly mean temperature, which was also best explained by a second-order polynomial.

From the fully parameterized model described above, we explored whether substituting monthly soil moisture, precipitation total, and mean temperature values with the corresponding values from the previous month improved model performance using Akaike information criterion (AIC) values. We retained soil moisture and precipitation totals from the months in which chickens were sampled. However, subsequent analyses included monthly mean temperature of the prior month because this time-lagged form of the temperature variable received stronger support than monthly mean temperature of the month of sampling (ΔAIC > 90). We used AIC scores to gauge whether polynomial terms for ALAN and mean temperature of the previous month improved model performance over linear effects of each. We also used AIC scores to arbitrate between spatial and non-spatial models and to evaluate the support for using an offset to account for the number of chickens sampled per site per month and to test whether there was any evidence for polynomial effects of predictor variables other than ALAN that reflected urbanization. We considered models with ΔAIC scores ≤ 2.0 as equally competitive and determined that a parameter estimate had a strong effect on WNV seroprevalence patterns when its 95% confidence interval (95% CI) did not overlap zero. Finally, we conducted two forms of model diagnostics. First, we checked for potential multicollinearity and redundancy among predictor variables with variance inflation factor (VIF) and considered VIF > 10 as potentially problematic sensu (61). Mean temperature of the previous month, and its quadratic term, were the only parameters with potentially problematic VIF scores. However, centering this variable at its mean resulted in VIF < 2.0 for each. Second, we assessed model performance with qqplots and residual vs. expected value plots using the simulateResiduals function in the R package *DHARMa* 0.2.4 (62).

## Author Statements

## Acknowledgements

We thank the Florida Department of Health for sharing their sentinel chicken surveillance data with us.

## Data Accessibility

Data will be made available via the Dryad repository.

